# HOOK2 downregulation compromises the tumorigenic and stemness properties of ovarian cancer cells by increasing endoplasmic reticulum stress

**DOI:** 10.1101/2025.05.29.656800

**Authors:** Elisa Suárez-Martínez, Sander R. Piersma, Thang V. Pham, Irene V. Bijnsdorp, Connie R. Jimenez, Amancio Carnero

## Abstract

Ovarian cancer stands out as one of the tumors with a high mortality rate in women. Therefore, the search for new therapeutic targets is essential to enhance patient prognosis. There is evidence suggesting that Hook2, an adaptor protein involved in microtubule transport, may play a role in cancer, particularly ovarian cancer. This study examines the role of HOOK2 in the progression of ovarian tumors. The findings reveal that a decrease in HOOK2 levels leads to diminished growth and cellular migration in ovarian cancer cells, impeding the *in vivo* formation of tumors. The reduction of HOOK2 is associated with both an increase in endoplasmic reticulum stress and an elevation in cell death, the latter likely caused by the activation of the unfolded protein response. Moreover, the study observes that the decrease in HOOK2 diminishes the properties of cancer stem cells in ovarian cancer, possibly due to the increase in cell death specifically found within these stem cells. Given the profound impact of reduced HOOK2 levels on ovarian cancer cells, this gene emerges as a promising therapeutic strategy for treating ovarian cancer patients.

## INTRODUCTION

Although ovarian cancer ranks as the 11th most common cancer among women, it stands out as the fifth leading cause of cancer-related death in women, proving to be the most fatal among all gyneacological tumors. With a 5-year relative survival rate remaining around only 50%, ovarian cancer poses a significant challenge (1, 2). The complexities in treating ovarian cancer are believed to arise from intratumoral heterogeneity, interactions with the microenvironment, and the presence of cancer stem cells (CSCs) in tumors. These stem cells exhibit heightened resistance to both radiotherapy and chemotherapy, contributing to relapses. For that reason, even though most patients initially respond well to the first line of therapy, the majority still experience recurrence. Subsequent responses to a second line of treatment tend to be only moderate (3). Consequently, it is essential to discover new targeted therapies to enhance the battle against ovarian cancer and address the frequent occurrence of relapses.

Previous studies have identified HOOK2 as a gene potentially associated with resistance to treatments in ovarian cancer patients (4). HOOK2 belongs to the family of HOOK dynein adapter proteins, involved in retrograde transport along microtubules (5–9). Each protein of this family is implicated in binding different organelles, facilitated by their variable C-terminal domain (10–13). Specifically, HOOK2 plays a crucial role in the composition, positioning, and functioning of centrosomes (12). Therefore, disruptions in this protein have been linked to the impairment of classic centrosomal functions, including division (14), polarized migration (15), and cilium formation (16). HOOK2 is also implicated in the positioning or formation of aggresomes—membrane-less cytoplasmic inclusions containing ubiquitinated and misfolded proteins, surrounded by intermediate filaments (17). Additionally, proteins in the HOOK family have been described to belong to the FHF complex, involved in various functions depending on its composition. Notably, the FHIP2A and HOOK2 complex has been observed to mediate the transport of intermediates from the endoplasmic reticulum to the Golgi apparatus (18, 19). The involvement of HOOK2 in cancer remains largely unexplored, and investigating its role in the initiation and maintenance of ovarian tumors could prove valuable in addressing the emergence of resistance to therapies for this specific tumor type.

Here we show that the reduction of HOOK2 inhibits the growth and migration of ovarian cancer cells, both *in vitro* and *in vivo*. The cells with downregulated HOOK2 increased endoplasmic reticulum (ER) stress, and concurrently exhibited an increase in cell death, likely as a consequence of the heightened ER stress, further intensified by the blockade of autophagic flux. Lastly, cells with diminished HOOK2 levels also demonstrated a decrease in CSC properties, potentially linked to the observed increase in cell death specifically within these stem cells.

## RESULTS

### 1. HOOK2 downregulation reduces cell growth and migration of ovarian cancer cells *in vitro* and impairs tumor formation *in vivo*

Analysis of patient databases of ovarian cancer revealed that approximately 14% of patients exhibit amplifications in this gene (**Figure 1A**), with these alterations seemingly linked to a decrease in overall survival (**Figure 1B**). Therefore, we decided to investigate the involvement of HOOK2 in these tumors by reducing its levels in two ovarian cancer cell lines (OVCAR-8 and SKOV-3). This reduction was achieved using the CRISPR-Cas9 technique in the OVCAR-8 line and through the use of shRNAs in the SKOV-3 line, due to the inability to generate CRISPR clones in the latter line. The downregulation of the gene was validated by Western Blot (WB) (**Figure 1C, S1A**), sequencing in the CRISPR clones (**Figure S1B**), and q-RT-PCR (**Figure S1C**). Cells with reduced levels of the target gene showed a decreased capacity for colony formation (**Figure 1D**) and migration (**Figure 1E**). Furthermore, in terms of growth, it was observed that the reduction of HOOK2 led to a decrease in *in vitro* proliferation (**Figure 1F**), especially in the SKOV-3 line, which became even more apparent *in vivo*. Cells with decreased levels of HOOK2 exhibited a reduction or complete inhibition of tumor growth when implanted in mouse models (**Figure 1G**). Only SKOV-3 cells carrying shA grew *in vivo,* probably due to loss of the shA construct during growth. In summary, HOOK2 is involved in maintaining key properties for tumor cells, such as cell proliferation or migration, and its reduction seems to negatively impact these cells.

**Figure 1.**
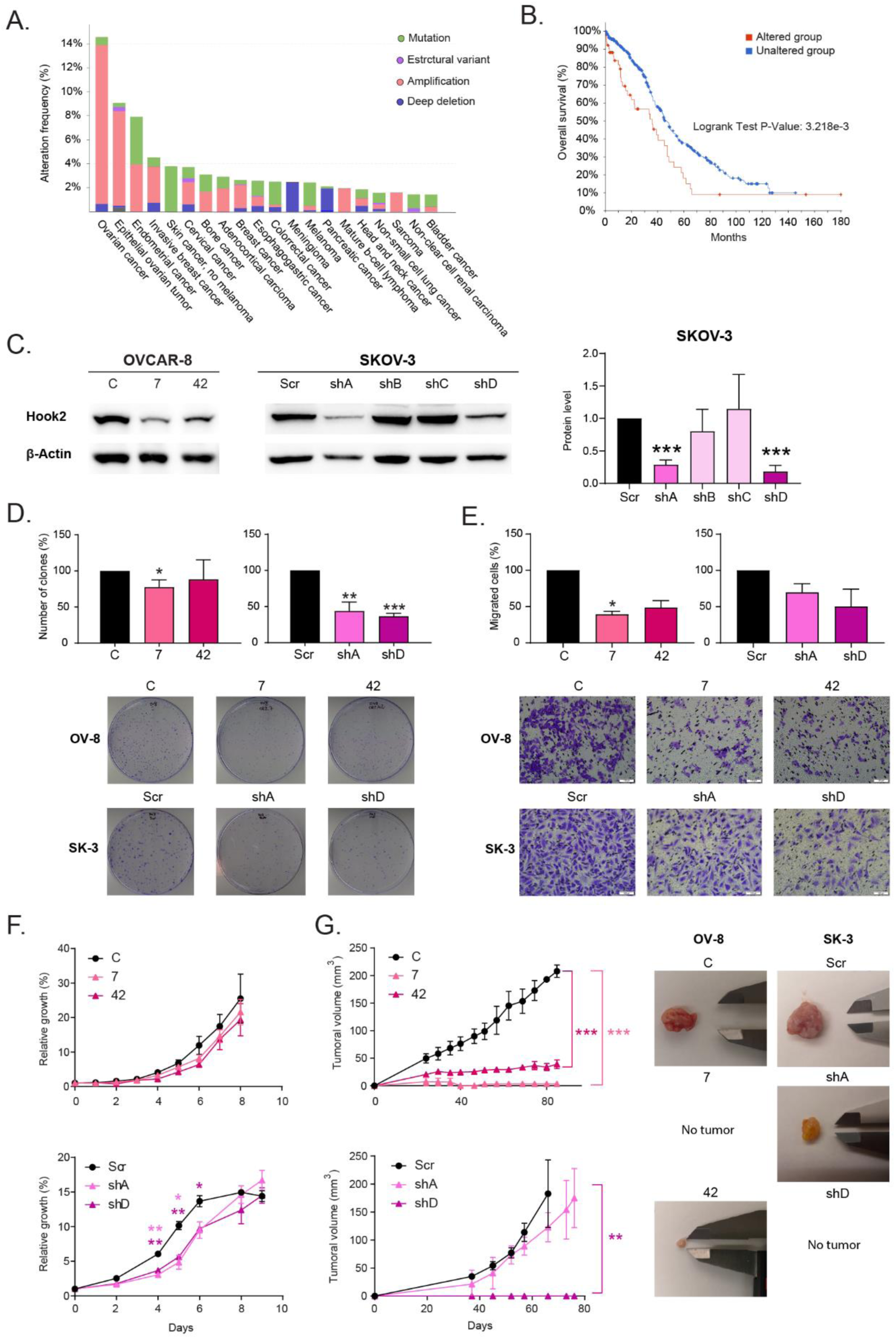
Hook2 is amplified in ovarian cancer patients, and its downregulation leads to a reduction in *in vitro* cell proliferation and migration, as well as *in vivo* tumor growth. **(A)** Analysis of genetic alterations in the HOOK2 gene across various tumor types. **(B)** Comparison of overall survival between patients with and without HOOK2 alterations. **(C)** Validation of decreased HOOK2 levels in ovarian cancer cell lines OVCAR-8 and SKOV-3 through WB. Assessment of the percentage of **(D)** clone formation, **(E)** migrating cells, and **(F)** relative growth of ovarian cancer cells with diminished HOOK2 *in vitro*. The mean of 3 independent experiments ± SEM is presented. **(G)** Evaluation of *in vivo* xenotransplant growth originating from cells with reduced HOOK2 levels compared to parental cells from OVCAR-8 and SKOV-3 cell lines. Graphs represent the tumor size (mean ± SEM). Representative images of the tumors are shown. Statistical analysis was performed with Student’s t test (*p < 0.05; **p < 0.01; ***p < 0.001). The lack of an asterisk indicates that the data do not reach statistical significance.

### 2. Reducing HOOK2 levels leads to an increase in endoplasmic reticulum stress in ovarian cancer cells, possibly due to its implication in aggresome formation

To investigate the underlying cause of the negative impact of reducing HOOK2 in ovarian cancer cells, a proteomic study was conducted, comparing two CRISPR clones of HOOK2 from the OVCAR-8 cell line with their parental counterparts. Approximately 6300 proteins were identified in this study (**Supplementary table 1**), revealing significant changes in expression of 164 proteins (p< 0.05) as a consequence of HOOK2 reduction (**Figure 2A, 2B, Supplementary table 2**). From this dataset, 111 proteins exhibiting most discriminatory alterations in expression (p< 0.05; logFC> 1/ logFC < −1) were subjected to gene ontology analysis to explore associated biological processes. Proteins that were upregulated following the reduction of HOOK2 were predominantly associated with protein folding, response to misfolded proteins, and more specifically, the regulation of ER stress (**Figure 2C, 2E, S2A, S2C, S2E**). In the case of the downregulated proteins, these were mainly involved in cell substrate junction assembly and, to a lesser extent, in the regulation of metabolic processes, such as peptidyl-proline modification and biosynthesis of modified amino acids, as well as translation regulation (**Figure 2D, S2B, S2D**).

**Figure 2.**
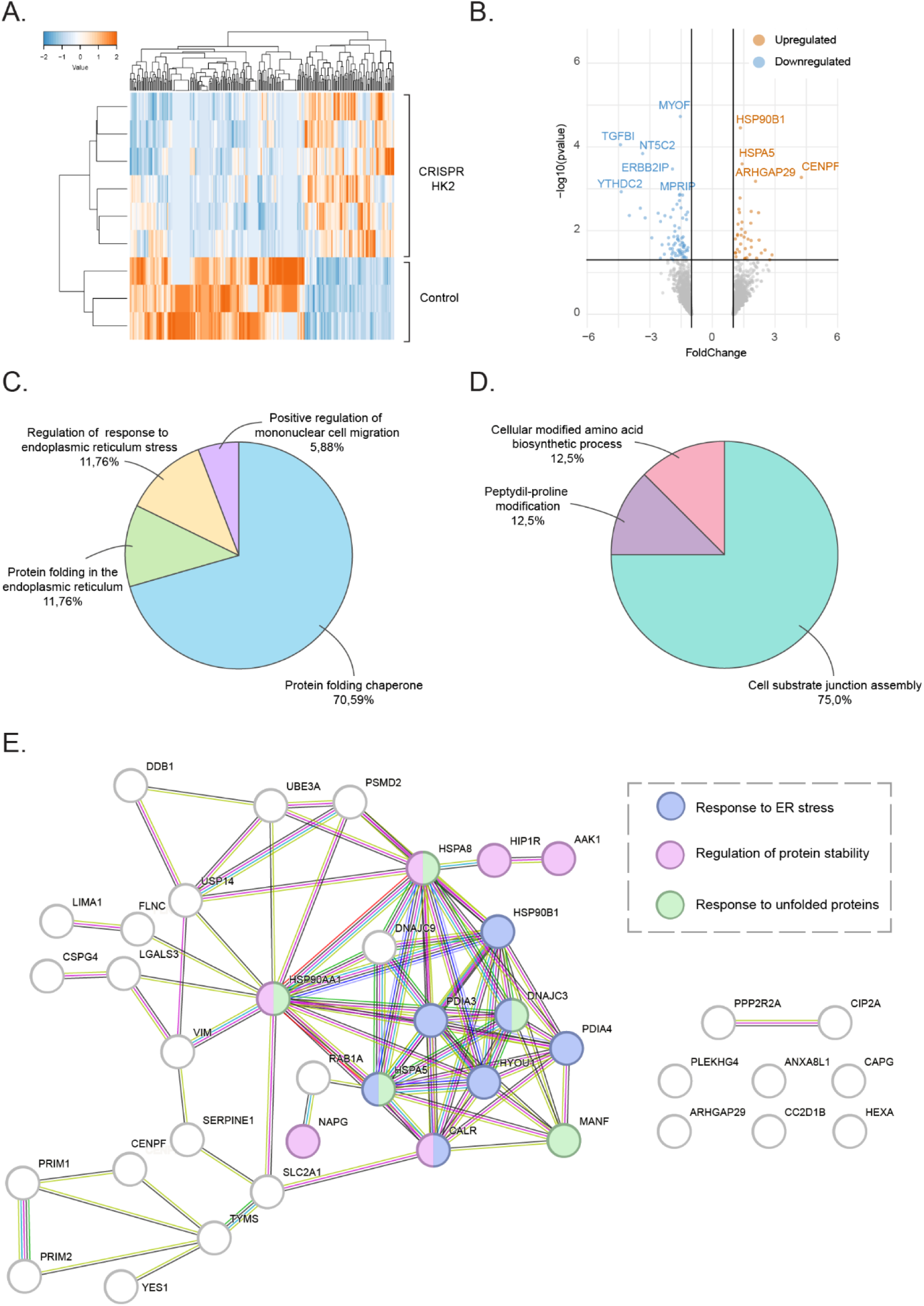
Proteomic analysis of cells exhibiting downregulated HOOK2 reveals significant alterations in processes associated with protein misfolding and ER stress. **(A)** Supervised clustering of proteins that either increase or decrease upon the reduction of HOOK2 expression in the ovarian cancer cell line OVCAR-8. **(B)** Volcano plot displaying the proteins that are upregulated or downregulated in OVCAR-8 cells with diminished HOOK2 levels. The names of the top 10 most significantly upregulated or downregulated proteins are showing. Ontology term analysis of proteins **(C)** upregulated or **(D)** downregulated when HOOK2 is reduced in the OVCAR-8 line. **(E)** Protein network of upregulated proteins upon HOOK2 reduction in OVCAR-8 cell line. Network nodes represent proteins and edges represent protein-protein associations. Some nodes are colored to show that they are associated to a specific GO term.

Based on the findings from the proteomic analysis, it seems that diminished levels of HOOK2 may be implicated in the unfolded protein response (UPR) within the ER. To validate this hypothesis, the expression of various markers associated with the UPR and the sensitivity of these cells to different ER stress inducers were assessed. The results indicated that the reduction of HOOK2 levels led to an upregulation of several UPR markers in ovarian cancer cells, such as ATF6 in the OVCAR-8 line, or ATF-4 and CHOP in the SKOV-3 line (**Figure 3A**). Furthermore, cells with decreased HOOK2 demonstrated heightened sensitivity to ER stress-inducing compounds, particularly Brefeldin A (**Figure 3B**). These findings confirm that the decrease in HOOK2 triggers the activation of the UPR in ovarian cancer cells, likely due to an accumulation of misfolded proteins.

**Figure 3.**
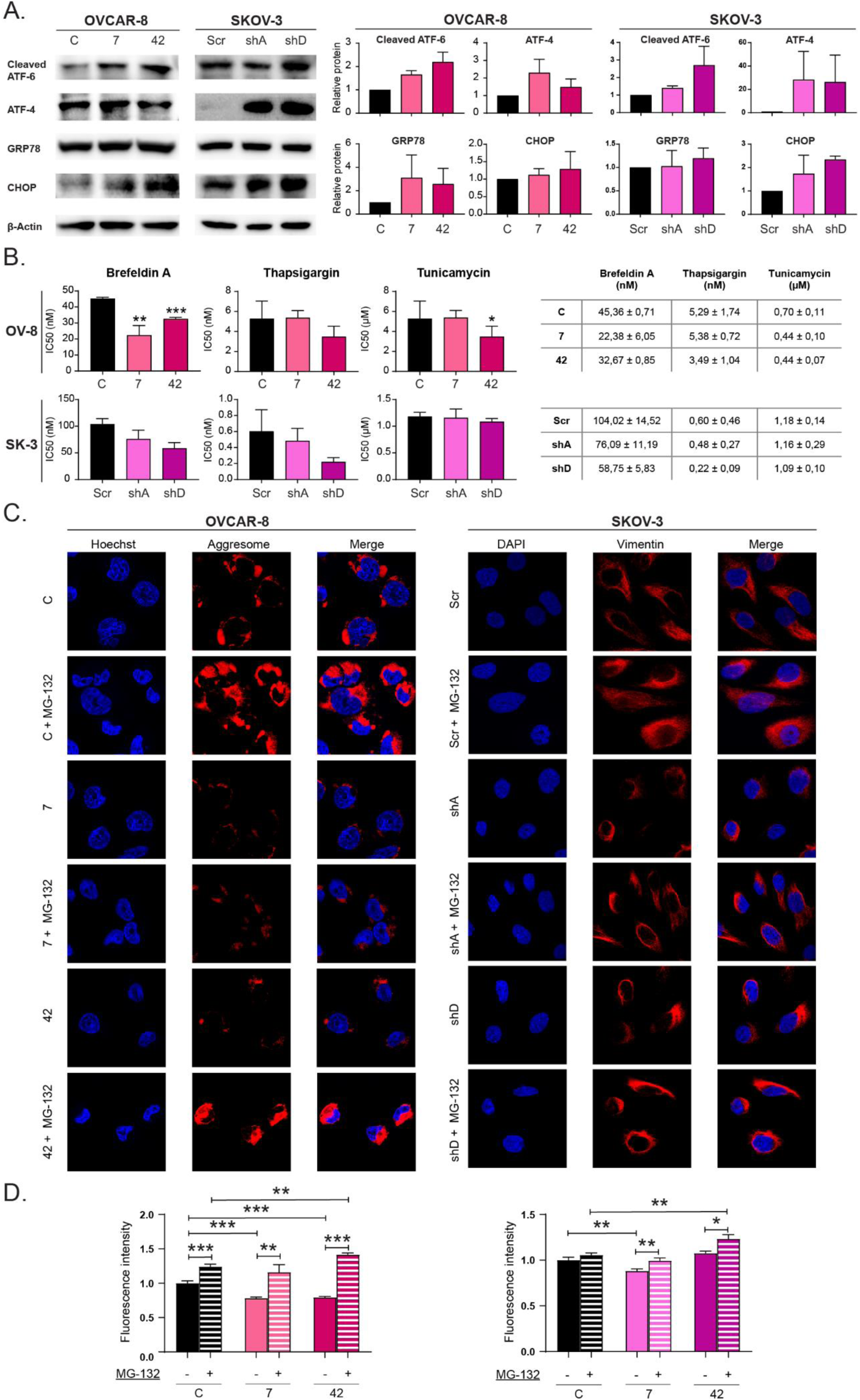
Cells with downregulated HOOK2 show heightened expression of ER stress markers, increased sensitivity to agents inducing this response, and difficulties in the formation of aggresomes. **(A)** WB analysis of proteins involved in the UPR in cells with reduced HOOK2 levels. **(B)** Measurement of the IC50 for drugs inducing ER stress in cells with decreased HOOK2. The mean of 3 independent experiments ± SEM is presented. Statistical analysis was performed with Student’s t test (*p < 0.05; **p < 0.01; ***p < 0.001). The lack of an asterisk indicates that the data do not reach statistical significance. **(C)** Immunofluorescence images depicting aggresomes (in red) in cells with reduced HOOK2 levels treated with the proteasome inhibitor MG-132. The nucleus is counterstained with DAPI (in blue), and the merged channels are displayed. **(D)** Quantification of aggresome marker fluorescence intensity normalized to untreated parental cells, using ImageJ software. Analysis involved a minimum of 100 cells for each condition, and statistical significance was assessed through Student’s t-test (*p<0.05; **p<0.01; ***p<0.001). The lack of an asterisk indicates that the data do not reach statistical significance.

While there has been no documented association between HOOK2 and ER stress to date, this protein has been linked to the formation of aggresomes, a structure involved in the degradation of misfolded proteins (17). We hypothesized that the reduction of HOOK2 might be triggering UPR activation due to a malfunction in aggresomes, leading to the accumulation of misfolded proteins. Thus, we examined aggresome formation in cells with downregulated HOOK2 in the presence and absence of the proteasome inhibitor MG-132. Proteasome inhibition is known to induce the accumulation of misfolded proteins in aggresomes and is commonly used as a positive control for aggresome analysis. Our findings revealed a reduction in aggresome formation under normal conditions in cells with reduced HOOK2 levels, particularly evident in the OVCAR-8 line. However, cells with downregulated HOOK2 treated with the proteasome inhibitor were able to recapitulate the aggresome formation observed in the parental lines (**Figure 3C-D**). Therefore, while the decrease in HOOK2 levels may impact aggresome formation, it is likely not the sole factor contributing to the increased ER stress observed in our study.

### 3. The decrease of HOOK2 causes an increased cell death in ovarian cancer cells that is dependent on the autophagic flux

We wondered whether the increase in ER stress observed upon HOOK2 reduction could explain the changes in the growth of ovarian cancer cells observed both *in vitro* and *in vivo*. Depite their initial protective function against proteotoxic stress, it is widely established that pathways activated in the UPR have the potential to induce cell death if sustained over time (20, 21). Therefore, we considered the possibility that cells with reduced levels of HOOK2 might be adversely affected by ER stress, leading to an increase in cell death. To investigate this, the percentage of apoptotic and necrotic cells, as well as the overall percentage of dead cells, was analyzed by flow cytometry upon reducing HOOK2. Additionally, certain markers of apoptotic cell death were examined through WB. While the reduction of HOOK2 showed a tendency to increase cell death in ovarian cancer cells, the effect was not statistically significant (**Figure 4A**). However, in the SKOV-3 line, an increase was observed in all studied apoptotic markers (**Figure 4B**). Therefore, cells with reduced levels of HOOK2 tend to exhibit a higher propensity for mortality.

**Figure 4.**
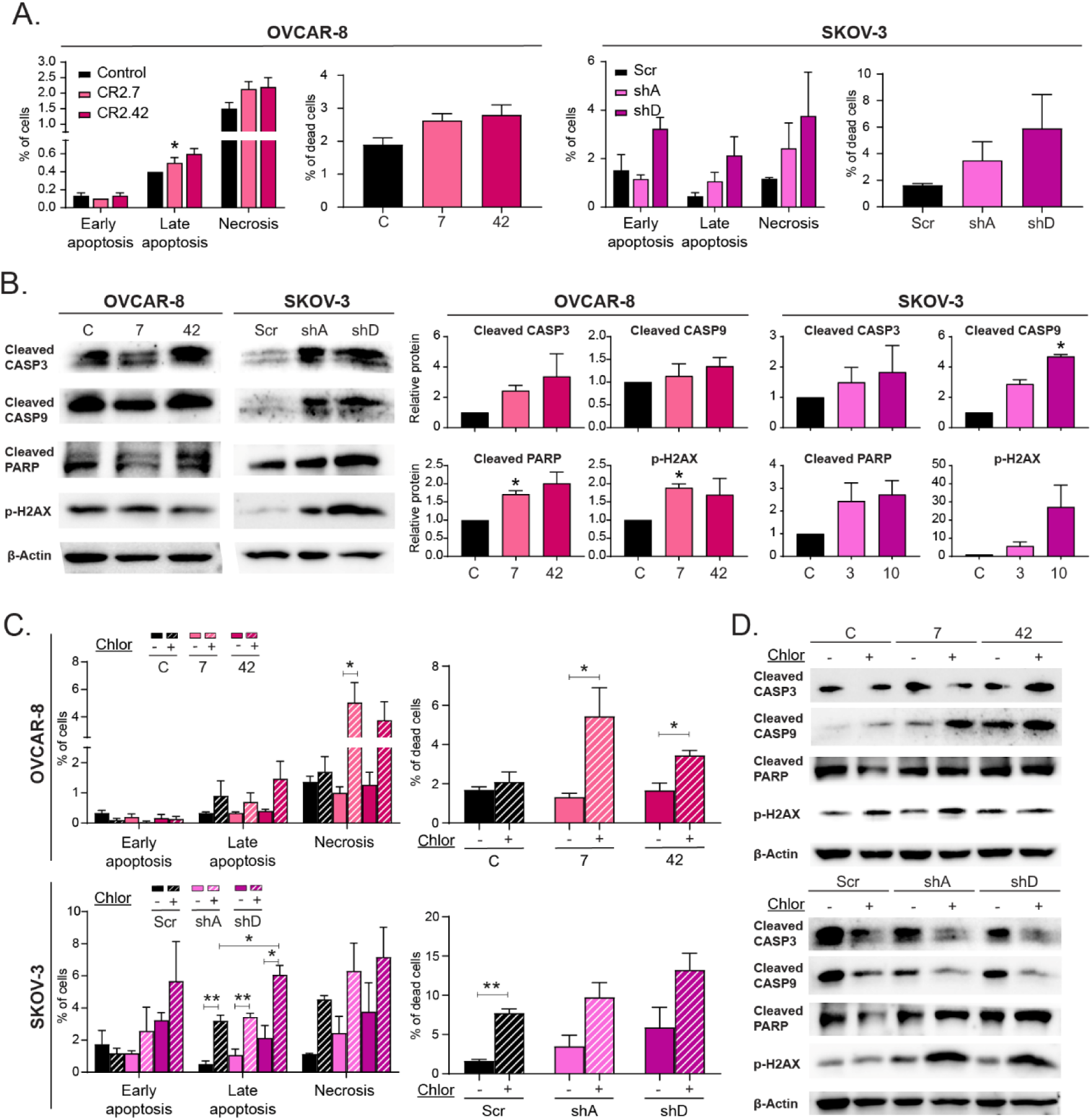
Decreased levels of HOOK2 may lead to increased cell death in ovarian cancer cells, which is dependent on autophagic flux. **(A)** Quantification by flow cytometry of the percentage of apoptotic and necrotic cells, as well as total cell death, when HOOK2 is downregulated. **(B)** WB analysis of proteins related to apoptotic cell death in cells with reduced HOOK2 expression. **(C)** Quantification by flow cytometry of the percentage of apoptotic and necrotic cells, as well as total cell death, in HOOK2-downregulated cells treated with the autophagic flux inhibitor chloroquine. The mean of 3 independent experiments ± SEM is presented. Statistical analysis was performed with Student’s t test (*p < 0.05; **p < 0.01; ***p < 0.001). The lack of an asterisk indicates that the data do not reach statistical significance. **(D)** Protein levels of apoptosis-associated proteins in HOOK2-downregulated cells treated with chloroquine.

On the other hand, it has been described that an increase in ER stress is capable of activating degradation via autophagy (22, 23). This mechanism would act compensatorily to degrade accumulations of misfolded proteins, preventing the prolonged stress from triggering cell death. Therefore, we hypothesized that, in our model, autophagy could be functioning as a protective mechanism against cell death. To validate this hypothesis, cells with reduced levels of HOOK2 were treated with chloroquine, an inhibitor of autophagic flux, and its impact on cell death was examined. It was observed that the inhibition of autophagic flux significantly affected cells with downregulated HOOK2 more than parental cells, leading to an increase in cell death of up to 15% (**Figure 4C-D**). Furthermore, cells appeared to undergo cell death through both apoptotic and non-apoptotic mechanisms, as confirmed by the increase in certain apoptotic markers observed when cells were treated with chloroquine (**Figure 4C-D**). These results indicate that autophagy acts as a pro-survival mechanism in ovarian cancer cells under conditions of elevated ER stress. In conclusion, the data suggest that cells with downregulated HOOK2 tend to exhibit higher mortality and are more sensitive to autophagic flux blockade.

### 4. CSCs properties are reduced in ovarian cancer cells by the downregulation of HOOK2, likely due to the increased cell death associated with a heightened ER stress

One of the underlying causes of treatment resistance and cancer relapses in ovarian cancer lies in the presence of CSCs within tumors. These cells exhibit increased resistance to radiotherapy, cytotoxic agents, and targeted therapies (24). Consequently, we questioned whether the reduction of HOOK2 might impact these CSCs, compromising their viability. To assess changes in CSC properties in cells with decreased HOOK2 levels, various analyses were conducted. Firstly, the proportion of holoclones, meroclones, and paraclones from the clonogenic assay was analyzed, revealing a significant reduction in the percentage of holoclones upon HOOK2 reduction (**Figure 5A**). Additionally, it was observed that cells with diminished HOOK2 levels formed fewer tumorspheres compared to parental cells, and these tumorspheres were also smaller in the OVCAR-8 line (**Figure 5B**). Lastly, the percentage of cells positive for the stem cell marker EpCAM was measured, indicating that, in all cases, cells with downregulated HOOK2 exhibited a diminished proportion of EpCAM-positive cells (**Figure 5C**). All these data suggest that the reduction of HOOK2 decreases the CSC properties of ovarian cancer cells.

**Figure 5.**
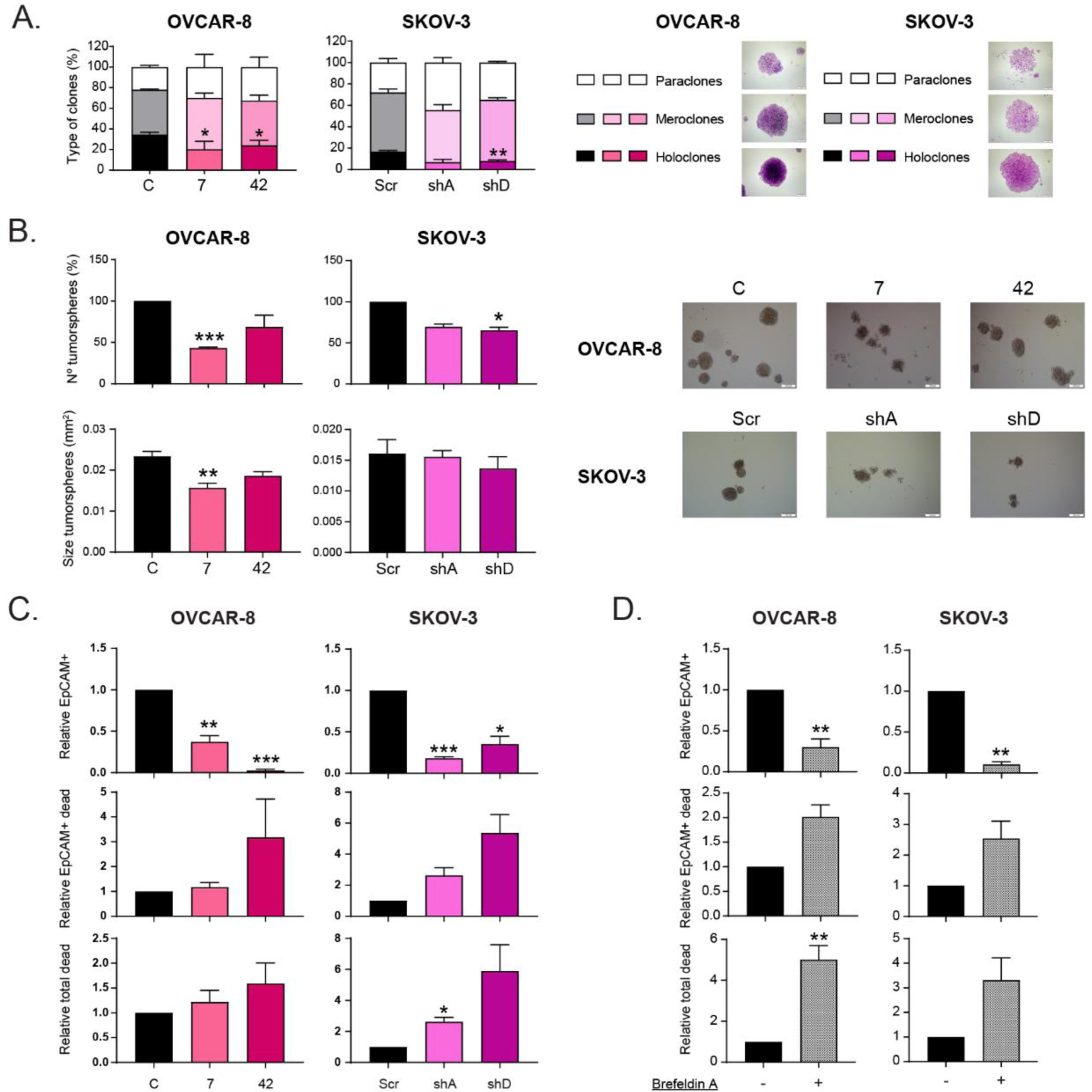
The properties of CSCs are impacted by the reduction of HOOK2, likely due to an increase in cell death associated with heightened ER stress. **(A)** Assessment of the percentage of holoclones, meroclones, and paraclones in a clonability assay upon decreasing HOOK2 levels. **(B)** Analysis of the number and size of tumorspheres generated from ovarian cancer cells with diminished HOOK2. **(C)** Quantification of the relative number of cells expressing the EpCAM marker, EpCAM-positive dead cells, and total dead cells in cells with reduced HOOK2. **(D)** Quantification of the relative number of cells expressing the EpCAM marker, EpCAM-positive dead cells, and total dead cells in parental ovarian cancer cells treated with the ER stress inducer brefeldin A. The mean of 3 independent experiments ± SEM is presented. Statistical analysis was performed with Student’s t test (*p < 0.05; **p < 0.01; ***p < 0.001). The lack of an asterisk indicates that the data do not reach statistical significance.

These alterations in stem cell properties could be attributed to the previously observed increase in cell death. To test the validity of this hypothesis, the mortality of EpCAM-positive cells with downregulated HOOK2 levels was investigated. It was noted that the reduction in HOOK2 levels led to an elevated percentage of cell death among EpCAM-positive cells (**Figure 5C**). Therefore, it is possible that the decline in CSC properties is indeed linked to a higher rate of cell death in this cell population when HOOK2 levels are reduced. Given our observation that this heightened cell death might be induced by increased ER stress, we explored whether an elevated level of ER stress alone could recapitulate the cell death observed in EpCAM-positive cells when HOOK2 is reduced. Treating parental cells with Brefeldin A to induce ER stress significantly reduced the percentage of EpCAM-positive cells and, furthermore, increased the percentage of dead EpCAM-positive cells (**Figure 5D**). Consequently, an increase in ER stress alone is capable of inducing a phenotype similar to that observed when HOOK2 is reduced. In conclusion, the reduction in HOOK2 results in a decline in CSC properties, likely due to an increased rate of cell death in these cells, possibly stemming from prolonged activation of the UPR.

## DISCUSION

The search for new therapies for ovarian cancer patients is essential to improving their prognosis. To this end, we investigated the role of HOOK2 in this type of tumor and explored whether its inhibition could represent a promising strategy. We showed that reducing HOOK2 reduces the growth and migration of ovarian cancer cells and leads to increased ER stress, accompanied by elevated cell death. This increased cell death is likely driven by the intensified ER stress and further aggravated by disrupting autophagic flux. Additionally, cells with reduced HOOK2 levels show a decline in CSC properties, which may be associated with the increased cell death observed specifically in these stem cell populations.

Regarding the reduction in growth and migration caused by the downregulation of HOOK2 in ovarian cancer cells (**Figure 1D, 1F, 1E**), while a direct relationship between HOOK2 and tumor cell migration has not been described, it has been noted that this gene is crucial for the binding of the PAR3/PAR6/aPKC complex with COMT. This complex plays a pivotal role in regulating polarized migration, and the decrease in HOOK2 levels has been observed to randomly redistribute the orientation of centrosomes during migration (15). This could provide a plausible explanation for the observed reduction of migration in tumor cells. Furthermore, the increased ER stress appreciated in these cells (**Figure 2C, 2E, and 3A-B**) could account for the observed reduction in, both, *in vitro* and *in vivo* growth. While a transient activation of the UPR can promote the survival of tumor cells (25), sustained activation of this mechanism over time may induce a cytotoxic effect, triggering activation of the apoptotic pathway and other modes of cell death, such as necroptosis, ferroptosis, or death through the iDISC platform (20, 21). In our study, we noted an increase, not only in apoptosis, but also in non-apoptotic forms of cell death upon downregulation of our gene of interest (**Figure 4**).

To date, no relationship has been described between HOOK2 and the occurrence of ER stress due to the accumulation of misfolded proteins. However, our data suggest that this could be attributed to HOOK2’s involvement in the formation of aggresomes (**Figure 3C-D**). Previous observations indicated that HOOK2 expression promotes the formation of these structures, and its dominant negative form is capable of inhibiting protein accumulation in this area (17). In our study, we observed that the reduction of HOOK2 hinders the formation of these structures in ovarian cancer cells (**Figure 3C-D**). Inhibition of aggresome formation has been linked to the accumulation of misfolded proteins and the onset of ER stress. For instance, the inhibition of histone deacetylase 6 (HDAC6), involved in the transport of misfolded proteins to aggresomes, can induce ER stress in tumor cells (26, 27). This phenomenon is associated with an increase in apoptosis in these cells and heightened sensitivity to the accumulation of misfolded proteins (26). Therefore, if the decrease in HOOK2 indeed interferes with aggresome formation, this could lead to the occurrence of ER stress in ovarian cancer cells.

On the other hand, although its role in aggresome formation may be crucial for the onset of ER stress, cells with downregulated HOOK2 treated with a proteasome inhibitor are still capable of forming aggresomes relatively normally (**Figure 3C-D**). Therefore, it is likely that other factors contribute to the induction of increased ER stress upon HOOK2 reduction. The

FTS/HOOK2/FHIP2A complex has been linked to the motility of tubular intermediaries that facilitate ER transport to the Golgi apparatus (19). Impaired transport between these two organelles can lead to elevated ER stress, similarly to what is observed with the drug Brefeldin A (28). Both processes are not mutually exclusive, and the reduction of HOOK2 may lead to a scenario where, not only aggresome formation decreases, but also the ER-Golgi transport is hindered, resulting in an even more significant increase in ER stress.

In cells with downregulated HOOK2, not only is there a decrease in growth and an increase in cell death, but also a reduction in properties associated with CSCs (**Figure 5A-C**). We speculate that this could be due to an increase in cell death specifically in CSCs. When analyzing cell death in EpCAM-positive cells, we find that cells with diminished HOOK2 levels exhibit higher mortality compared to parental cells (**Figure 5C**). This result supports the hypothesis that the decrease in HOOK2 not only leads to an overall decrease in the survival of ovarian cancer cells, but also specifically impacts CSCs. The diminished viability of CSCs could be linked to the observed increase in ER stress resulting from HOOK2 reduction. Numerous studies have demonstrated that maintaining proteostasis by the ER is crucial for preserving the integrity of CSCs. Inducing the UPR has been shown to diminish CSC properties in various cancers, including colorectal carcinoma (29–31), breast cancer (32), head and neck cancer (33), glioblastoma (34, 35), and prostate carcinoma (36). Therefore, the heightened ER stress could contribute to the observed reduction in CSCs capabilities in our model with reduced HOOK2. In fact, we have observed that pharmacological induction of ER stress in ovarian cancer cells recapitulates the phenotype of decreased HOOK2 in terms of CSC mortality (**Figure 5D**). Hence, these findings strengthen the hypothesis that the decrease in the gene of interest leads to an increase in CSC mortality due to the rise in ER stress.

Because the reduction of HOOK2 compromises the viability of ovarian cancer cells and impedes tumor formation *in vivo*, the use of inhibitors targeting this protein could be considered a promising therapeutic strategy for ovarian cancer patients. Drugs capable of blocking aggresome formation (27, 37) or inducing ER stress (38–41) have been employed in various studies to treat different types of tumors. In many instances, these drugs have been used in combination with clinically approved chemotherapeutics, paired with immunotherapeutic agents, or alongside other drugs capable of inducing proteotoxic stress, such as proteasome inhibitors. Considering the successful use of these drugs in cancer, HOOK2 could be a valuable therapeutic target, given that the reduction of this protein can initiate proteotoxic stress in tumor cells. Therefore, it would be interesting to explore the potential inhibition of HOOK2 as a treatment for ovarian cancer, both as a standalone therapy and in combination with chemotherapeutic agents or proteotoxic stress-inducing drugs.

## METHODS

### Cell culture

SKOV-3 and OVCAR-8 cell lines were obtained from the ATCC commercial repository and maintained in RPMI (AQmedia: Sigma) with 10% fetal bovine serum (FBS) (Gibco), penicillin, streptomycin and fungizone (Sigma) and incubated at 37 °C in 5% CO2 in a humidified atmosphere. Cells were negative for mycoplasma.

### CRISPR/Cas9 generation of HOOK2 knockdown

To establish knockdown models targeting the HOOK2 sequence (exon 11), a specific sgRNA with the sequence CGCCGAGCGCACGCGACAAC was employed. First, cells were infected with a virus containing the sgRNA, followed by selection with puromycin (1 µg/ml). Subsequently, cells were isolated using single-cell sorting via the FACS Jazz system (BD Biosciences) into 96-well plates. After one month, samples from wells displaying growth were amplified and subjected to validation through Western blot analysis. The selected CRISPRs were sequenced by the Genomics and Sequencing service at IBiS.

### Clonogenic assay and clonal heterogeneity analysis

To evaluate the cellular capacity to form individual clones, triplicates of 1×10^4 cells were plated in 10 cm plates. Cells were fixed with 0.5% glutaraldehyde and stained with 0.5% crystal violet after 10 days. The number of clones was counted, and the types of clones were classified according to phenotype and ability to reconstitute the culture.

### Migration assay (Boyden chamber)

A total of 9×10^5 cells (OVCAR-8) or 4×10^5 cells (SKOV-3) were suspended in FBS-free medium and subsequently seeded into an 8 μm Boyden chamber (Transwell). The chamber was placed in a 24-well plate containing FBS medium. After 24 hours, the cells were fixed using 0.5% glutaraldehyde and stained with 0.5% crystal violet. To eliminate nonmigrated cells, the inner membrane of the chamber was carefully cleaned. Then, images were captured at 20x magnification, and the migrated cells were quantified using an inverted microscope (Olympus IX-71).

### Growth curve

To assess proliferative capacity, a total of 5×10^4 cells were seeded in triplicate within 12-well plates. After 24 hours (Day 0), cells were fixed using 0.5% glutaraldehyde (Sigma), and subsequently, every 24/48 hours, a data point was captured over a span of 9 days. Following data collection, the plates were stained with 0.5% crystal violet (Sigma). The crystal violet stain was then solubilized in 20% acetic acid (Sigma) and quantified by measuring the absorbance at 595 nm, serving as a relative indicator of cell number. All values are expressed relative to the initial measurement on Day 0.

### Xenograft in nude mice

Tumorigenicity was assayed by the subcutaneous injection of 4×106 (OVCAR-8) or 5×106 (SKOV-3) cells into the right flanks of 4-week-old female athymic nude mice. Cells were suspended in Matrigel (Corning) prior to the injection. Animals were examined weekly. After 80-120 days, depending on the cell lines, mice were sacrificed, and tumors were extracted and conserved at −80 °C. Tumor volume (mm3) was measured using calipers. All animal experiments were performed according to the experimental protocol approved by the IBIS and HUVR Institutional Animal Care and Use Committee (0309-N-15).

### Cytotoxicity assay

In triplicate, a total of 1.2×10^4 cells per well were seeded in a 96-well plate. The following day, the cells were treated with decreasing concentrations of the compounds brefeldin A (10-0 μM), thapsigargin (1-0 μM) and tunicamycin (100-0 μM). After 96 h, the cells were stained with 0.5% crystal violet. Then, crystal violet was solubilized in 20% acetic acid (Sigma) and quantified at 595 nm absorbance to measure cell viability.

### Tumorsphere assay

A total of 3×104 cells were seeded in triplicate in 24-well Ultra-Low Attachment Plates (Costar) containing 1 mL of MammoCult basal medium (Stem Cell Technologies) supplied with 10% MammoCult proliferative supplement, 4 μg/mL heparin, 0.48 μg/mL hydrocortisone, penicillin and streptomycin. After 7 days, the number of primary tumorspheres formed was measured using an inverted microscope (Olympus IX-71).

### Cell death assay

The cells were maintained in culture with or without chloroquine for 48-72 h (OVCAR-8 [25 μm], SKOV-3 [50 μM]). Then, both suspended cells and adherent cells were collected, and the ‘Apoptosis Detection’ kit (Immunostep) was used to measure cell death following the manufacturer’s instructions. A FACSCanto II flow cytometer (BD Biosciences) was employed to detect the staining, and the results were analyzed using Diva software.

### Cell death assay in cells labelled with EpCAM by flow cytometry

First, a previously trypsinized suspension of 1×106 cells was resuspended in PBS with 2% FBS and 5 mM EDTA. Next, the cells were incubated with blocking agent (Miltenyi Biotec) for 10 minutes at 4 °C, and then, antibody labeling (Miltenyi Biotec) was performed for 15 minutes at room temperature and 15 minutes at 4 °C. Following this, two washes were performed using PBS with 2% FBS and 5 mM EDTA. After that, the previously described kit (Inmunostep) was used to label dead cells. Finally, the cells were examined using a FACSCanto II flow cytometer (BD Biosciences), and the results were analyzed using Diva software.

### RT‒qPCR

Total RNA from cell lines was extracted and purified using the ReliaPrepTM RNA Tissue Miniprep System (Promega), and reverse transcription was performed using the High Capacity cDNA Reverse Transcription kit (Life Technologies) according to the manufacturer’s instructions. The qPCR mixture contained the reverse transcriptase reaction product (16,6 ng/μl), 2.5 µL of water, 5 µL of GoTaqR Probe qPCR Master Mix (Promega) and 0.5 µL of the appropriate TaqMan Assay (IDT). The following probes were used: HPRT1 (Hs.PT.58.v.45621572) as an endogenous control and HOOK2 (Hs.PT.58.1901417).

### Protein isolation and Western blot analysis

Western blotting was performed according to standard procedures. We used the following primary antibodies: anti-HOOK2 (Abcam, ab151756), anti-ATF6α (Santa Cruz Biotechnology, sc-166659), anti-ATF4 (Santa Cruz Biotechnology, sc-390063), anti-GRP78 (Santa Cruz Biotechnology, sc-13539), anti-CHOP (Cell Signaling, #2895), anti-CAP3 (Cell Signaling, #9664), anti-CASP9 (Cell Signaling, #9502), anti-PARP (Cell Signaling, #9532), anti-p-H2AX (Ser139) (Cell Signaling, #9718), anti-LC3B (Abcam, ab48394), anti-p62 (Abcam, ab109012), and anti-B-actin (Abcam, ab16039) as a loading control. We used the following secondary antibodies: rabbit anti-mouse (Abcam, ab97046) and goat anti-rabbit (Abcam, ab97051). The proteins were detected using an ECL detection system (Amersham Biosciences) and a Bio-Rad Chemidoc Touch.

### Proteomic analysis

#### Gel electrophoresis and in-gel digestion of proteins

Gel electrophoresis and in-gel digestion were performed as described before (42). Briefly, protein lysates were separated on precast 4–12% gradient gels using the NuPAGE SDS-PAGE system (Invitrogen, Carlsbad, CA). Following electrophoresis, gels were fixed in 50% ethanol/3% phosphoric acid solution and stained with Coomassie R-250. Subsequently, the gels were washed, and proteins reduced and alkylated by incubating the whole gel in dithiothreitol and iodoacetamide, respectively. Gel lanes were cut into 5 bands and each band was cut into ∼1 mm3 cubes. The gel cubes from one band were transferred into an eppendorf tube and incubated with trypsin o/n. The peptides from each gel band were extracted and stored at −20 °C until LC-MS/MS analysis.

#### LC-MS/MS

Peptides were separated by an Ultimate 3000 nanoLC-MS/MS system (Dionex LC-Packings) equipped with a 45 cm × 75 μm ID fused silica column custom packed with 1.9 μm 120 Å ReproSil Pur C18 aqua (Dr Maisch GMBH). After injection, peptides were trapped at 6 μl/min on a 10 mm ×100 μm ID trap column packed with 5 μm 120 Å ReproSil Pur C18 aqua in 0.05% formic acid. Peptides were separated at 300 nl/min in a 10–40% gradient (buffer A: 0.5% acetic acid (Fisher Scientific), buffer B: 80% ACN, 0.5% acetic acid) in 60 min (100-min inject-to-inject). Eluting peptides were ionized at a potential of +2 kVa into a Q Exactive mass spectrometer (Thermo Fisher). Intact masses were measured at resolution 70,000 (at m/z 200) in the orbitrap using an AGC target value of 3E6 charges. The top 10 peptide signals (charge-states 2+ and higher) were submitted to MS/MS in the HCD (higher-energy collision) cell (1.6 amu isolation width, 25% normalized collision energy). MS/MS spectra were acquired at resolution 17,500 (at m/z 200) in the orbitrap using an AGC target value of 1E6 charges, a maxIT of 60 ms, and an underfill ratio of 0.1%. Dynamic exclusion was applied with a repeat count of 1 and an exclusion time of 30 s.

#### Protein identification and label-free quantitation

MS/MS spectra were searched against the reference proteome FASTA file (42161 entries; swissprot_2017_03_human_canonical_and_isoform). Enzyme specificity was set to trypsin, and up to two missed cleavages were allowed. Cysteine carboxamidomethylation (Cys, +57.021464 Da) was treated as fixed modification and methionine oxidation (Met, +15.994915 Da) and N-terminal acetylation (N-terminal, +42.010565 Da) as variable modifications. Peptide precursor ions were searched with a maximum mass deviation of 4.5 ppm and fragment ions with a maximum mass deviation of 20 ppm. Peptide and protein identifications were filtered at an FDR of 1% using the decoy database strategy. The minimal peptide length was seven amino acids. Proteins that could not be differentiated based on MS/MS spectra alone were grouped into protein groups (default MaxQuant settings). Searches were performed with the label-free quantification option selected. Proteins were quantified by spectral counting, that is, the number of identified MS/MS spectra for a given protein (43) combining the five fractions per sample. Raw counts were normalized on the sum of spectral counts for all identified proteins in a particular sample, relative to the average sample sum determined with all samples. To find statistically significant differences in normalized counts between sample groups, we applied the beta-binomial test (44), which takes into account within-sample and between-sample variation using an alpha level of 0.05. The generated data were filtered using the R platform to obtain proteins whose expression increased (FC > 1) or decreased (FC < −1) significantly (p < 0.05) and analyzed using the ClueGO app for Cytoscape (45) and ShinyGO 0.80 (http://bioinformatics.sdstate.edu/go/) and STRING (https://string-db.org/) platforms.

### Autophagy analysis by immunofluorescence

Cells were seeded onto glass coverslips and grown with or without chloroquine for 24 h (OVCAR-8 [25 μm], SKOV-3 [50 μM]). Then, they were fixed with 4% paraformaldehyde for 20 min and permeabilized with 0.5% Triton X-100 for 5 min. The coverslips were incubated with blocking solution (PBS + 0.1% Triton X-100 + 3% BSA) for 1 h and then incubated with anti-LAMP2 antibody (1:250, Abcam, ab25631) overnight at 4 °C. The coverslips were washed four times with PBS + 0.1% Triton X-100 and incubated overnight at 4 °C with the second primary antibody, anti-LC3B (1:250, Abcam, ab48394). The secondary antibodies anti-mouse Alexa Fluor 488 (1:250, Thermo Fisher A-11029) and anti-rabbit Alexa Fluor 633 (1:250, Thermo Fisher A-21071) were used. The nuclei were stained with DAPI, and the coverslips were mounted with ProLong Gold Antifade (Life Technologies). A confocal ultraspectral microscope (Leica Stellaris 8) that allowed sequential scanning of emission channels was used for image detection, and the images were analyzed with *ImageJ/Fiji* software.

### Aggresome detection

Cells were seeded onto glass coverslips and grown with or without MG-132 for 24 h (OVCAR-8 [0,35 μm], SKOV-3 [1 μM]). Then, they were fixed with 4% paraformaldehyde for 20 minutes. Staining of misfolded and accumulated proteins in the aggresome was performed in two different ways. Either, using immunofluorescence staining as described in the previous section with the primary antibody anti-Vimentin (1:250, Santa Cruz Biotechnology, sc-6260), or using the “PROTEOSTAT® Aggresome Detection Kit” (Enzo), following the manufacturer’s instructions. Briefly, for the use of this kit, cells were incubated for 30 minutes with the permeabilization solution (0.5% Triton X-100, 3 mm EDTA pH 8.0, 1x Assay Buffer) on ice and with agitation. Subsequently, 2 washes of 5 minutes each were performed with PBS at room temperature with agitation. It was then incubated for another 30 minutes with “Dual detection reagent” (1x Assay Buffer + PROTEOSTAT + Hoechst) at the dilution indicated in the kit. Two additional washes with PBS were performed, and the crystals were mounted on slides. Image capture and analysis were carried out like in the previous section.

### Statistical analysis

Statistical analyses for the experiments were performed using GraphPad Prism. Comparisons between control samples, CRISPR clones, and shRNAs were conducted using either unpaired Student’s t-test or Student’s t-test with Welch’s correction, as deemed appropriate. Each experiment was independently performed a minimum of three times, with triplicate samples in each instance. Statistical significance was defined as p-values less than 0.05 and represented using the following classification: p < 0.05 (*), p < 0.01 (**), and p < 0.001 (***).

## ABBREVIATIONS

Chlor: chloroquine
CSCs: cancer stem cells
ER: endoplasmic reticulum
FC: fold-change
IC50: half-maximal inhibitory concentration
RT-qPCR: Reverse transcription-quantitative PCR
sgRNA: single guide-RNA
UPR: unfolded protein response
WB: western blot

## DECLARATIONS

### ETHICS APPROVAL AND CONSENT TO PARTICIPATE

All methods were performed in accordance with the relevant guidelines and regulations of the Institute for Biomedical Research of Seville (IBIS) and University Hospital Virgen del Rocio (HUVR). All animal experiments and the entire procedure of the patient cohort were performed according to the experimental protocol approved by HUVR Animal Ethics (CEI 0309-N-15).

### CONSENT FOR PUBLICATION

Not applicable.

### AVAILABILITY OF DATA AND MATERIAL

All data and material will be available upon reasonable request. Raw proteomics data can be downloaded from the ProteomeXchange repository (dataset identifier PXD050782).

### COMPETING INTERESTS

The authors declare that they have no competing interests.

### FUNDING

This research was supported by grants from Ministerio de Ciencia, Innovación y Universidades (MCIU) Plan Estatal de I+D+I 2018, Agencia Estatal de Investigación (AEI) and Fondo Europeo de Desarrollo Regional (MCIU/AEI/FEDER, UE): RTI2018-097455-B-I00 and PID2021-122629OB-I00 funded by MCIN/AEI/10.13039/501100011033 and by “ERDF A way of making Europe”, by the “European Union”; AEI-MICIU/FEDER (iDIFFER network RED2022-134792-T). Additional grants from CIBER de Cáncer (CB16/12/00275), from Consejeria de Salud (PI-0397-2017) and Project P18-RT-2501 from 2018 competitive research projects call within the scope of PAIDI 2020—80% co-financed by the European Regional Development Fund (ERDF) from the Regional Ministry of Economic Transformation, Industry, Knowledge and Universities. Junta de Andalucía. Special thanks to the AECC (Spanish Association of Cancer Research) Founding Ref. GC16173720CARR for supporting this work.

## ACKNOWLEDGEMENTS

Special thanks to the AECC Foundation for supporting this work.

## AUTHORS CONTRIBUTION

AC conceived and designed this study. ES-M, SRP (proteomic) and IVB (proteomic) performed the experiments; AC and CRJ supervised the experiments. ES-M, SRP (proteomic), TVP (proteomic), CRJ (proteomic) and AC analyzed and interpreted the data. ES-M and AC drafted and edited the manuscript. All authors revised the manuscript.

## Corresponding author

Correspondence to Amancio Carnero.

## AUTHOR INFORMATION

### Authors and Affiliations

**Elisa Suarez-Martinez & Amancio Carnero.**

Instituto de Biomedicina de Sevilla (IBIS)/HUVR/CSIC, Hospital Universitario Virgen del Rocío, Ed. IBIS, Consejo Superior de Investigaciones Científicas, Universidad de Sevilla, Avda. Manuel Siurot S/N, 41013, Seville, Spain.

CIBER de Cancer (CIBERONC), Instituto de Salud Carlos III, Madrid, Spain.

**Sander R Piersma, Thang V Pham, Irene V Bijnsdorp & Connie R. Jimenez.**

OncoProteomics Laboratory, Dept. Medical Oncology, Amsterdam University Medical Center. De Boelelaan 1117, 1081HV Amsterdam, The Netherlands.

## Notes

https://proteomecentral.proteomexchange.org/cgi/GetDataset?ID=PXD050782

### Competing Interest Statement

The authors have declared no competing interest.

