## Supplementary Figures for "HOOK2 downregulation compromises the tumorigenic and stemness properties of ovarian cancer cells by increasing endoplasmic reticulum stress"

### SUPPLEMENTARY FIGURE 1

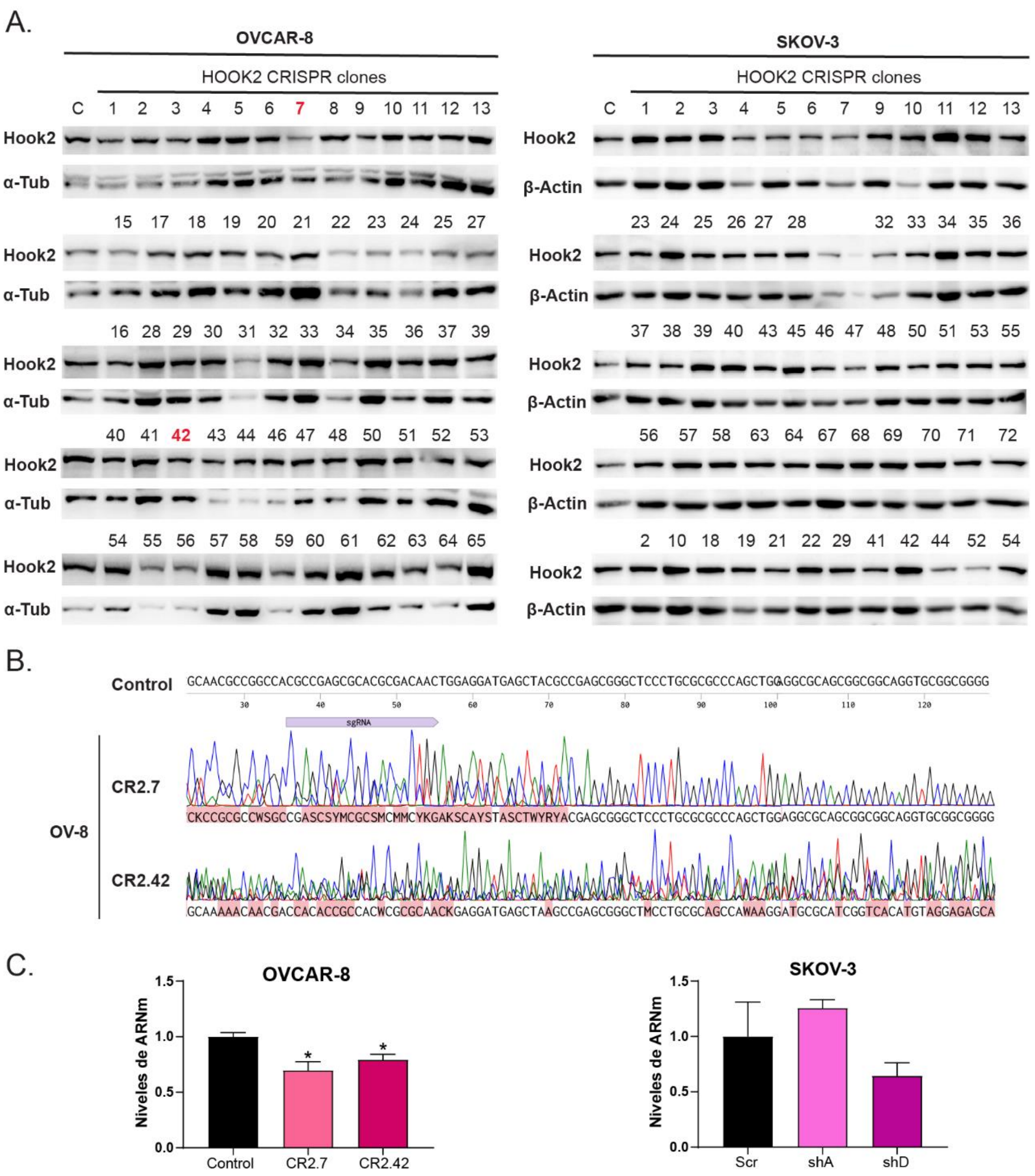

### SUPPLEMENTARY FIGURE 2

A.

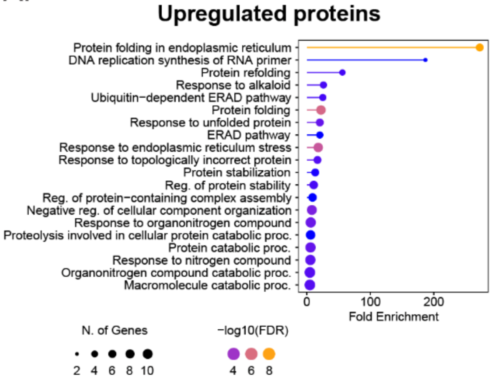


B.

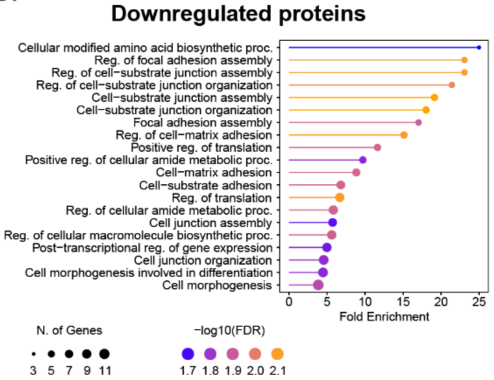


### C. Upregulated proteins

| Process | Terms | Associated Proteins Found |
|---|---|---|
| Positive regulation of mononuclear cell migration | Positive regulation of mononuclear cell migration (GO:0071677) | CALR, LGALS3, SERPINE1 |
| Regulation of response to endoplasmic reticulum stress | Negative regulation of response to endoplasmic reticulum stress (GO:1903573) | HSPA1A, HSPA5, HYOU1, USP14 |
| | Regulation of response to endoplasmic reticulum stress (GO:1905897) | HSPA1A, HSPA5, HYOU1, MANF, USP14 |
| Protein folding in the endoplasmic reticulum | Protein folding in endoplasmic reticulum (GO:0034975) | CALR, DNAJC3, HSP90B1, HSPA5, PDIA3 |
| | Ubiquitin-dependent ERAD pathway (GO:0030433) | CALR, HSP90B1, HSPA5, USP14 |
| Protein folding chaperone | 'De novo' protein folding (GO:0006458) | HSPA1A, HSPA1B, HSPA5, HSPA8 |
| | Protein refolding (GO:0042026) | HSP90AA1, HSPA1A, HSPA1B, HSPA5, HSPA8 |
| | Protein folding chaperone (GO:0044183) | CALR, HSP90AA1, HSPA1A, HSPA1B, HSPA5, HSPA8 |
| | Chaperone-mediated protein folding (GO:0061077) | HSPA1A, HSPA1B, HSPA5, HSPA8, PDIA4 |
| | Response to topologically incorrect protein (GO:0035966) | DNAJC3, HSP90AA1, HSPA1A, HSPA1B, HSPA5, HSPA8, MANF |
| | 'De novo' posttranslational protein folding (GO:0051084) | HSPA1A, HSPA1B, HSPA5, HSPA8 |
| | Response to unfolded protein (GO:0006986) | DNAJC3, HSP90AA1, HSPA1A, HSPA1B, HSPA5, HSPA8, MANF |
| | Interleukin-8 production (GO:0032637) | HSPA1A, HSPA1B, RAB1A, SERPINE1 |
| | Cellular response to heat (GO:0034605) | HSP90AA1, HSPA1A, HSPA1B |
| | Chaperone cofactor-dependent protein refolding (GO:0051085) | HSPA1A, HSPA1B, HSPA5, HSPA8 |
| | Regulation of interleukin-8 production (GO:0032677) | HSPA1A, HSPA1B, RAB1A, SERPINE1 |
| | Positive regulation of interleukin-8 production (GO:0032757) | HSPA1A, HSPA1B, RAB1A, SERPINE1 |

### D. Downregulated proteins

| Process | Term | Associated Genes Found |
|---|---|---|
| Peptidyl-proline modification | Peptidyl-proline modification (GO:0018208) | FKBP4, FKBP5, P4HA1 |
| Cellular modified amino acid biosynthetic process | Cellular modified amino acid biosynthetic process (GO:0042398) | ALDH7A1, CKB, GAMT |
| Cell substrate junction assembly | Cell-substrate junction organization (GO:0150115) | FERMT2, LAMC1, SLK, THBS1, VCL |
| | Regulation of cell-substrate junction organization (GO:0150116) | FERMT2, SLK, THBS1, VCL |
| | Cell-substrate junction assembly (GO:0007044) | FERMT2, LAMC1, SLK, THBS1, VCL |
| | Focal adhesion assembly (GO:0048041) | FERMT2, SLK, THBS1, VCL |
| | Regulation of cell-substrate junction assembly (GO:0090109) | FERMT2, SLK, THBS1, VCL |
| | Regulation of focal adhesion assembly (GO:0051893) | FERMT2, SLK, THBS1, VCL |

E.

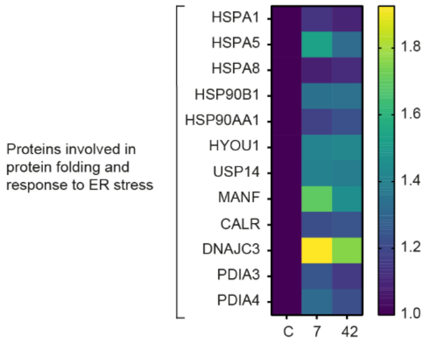

### SUPPLEMENTARY FIGURE 3

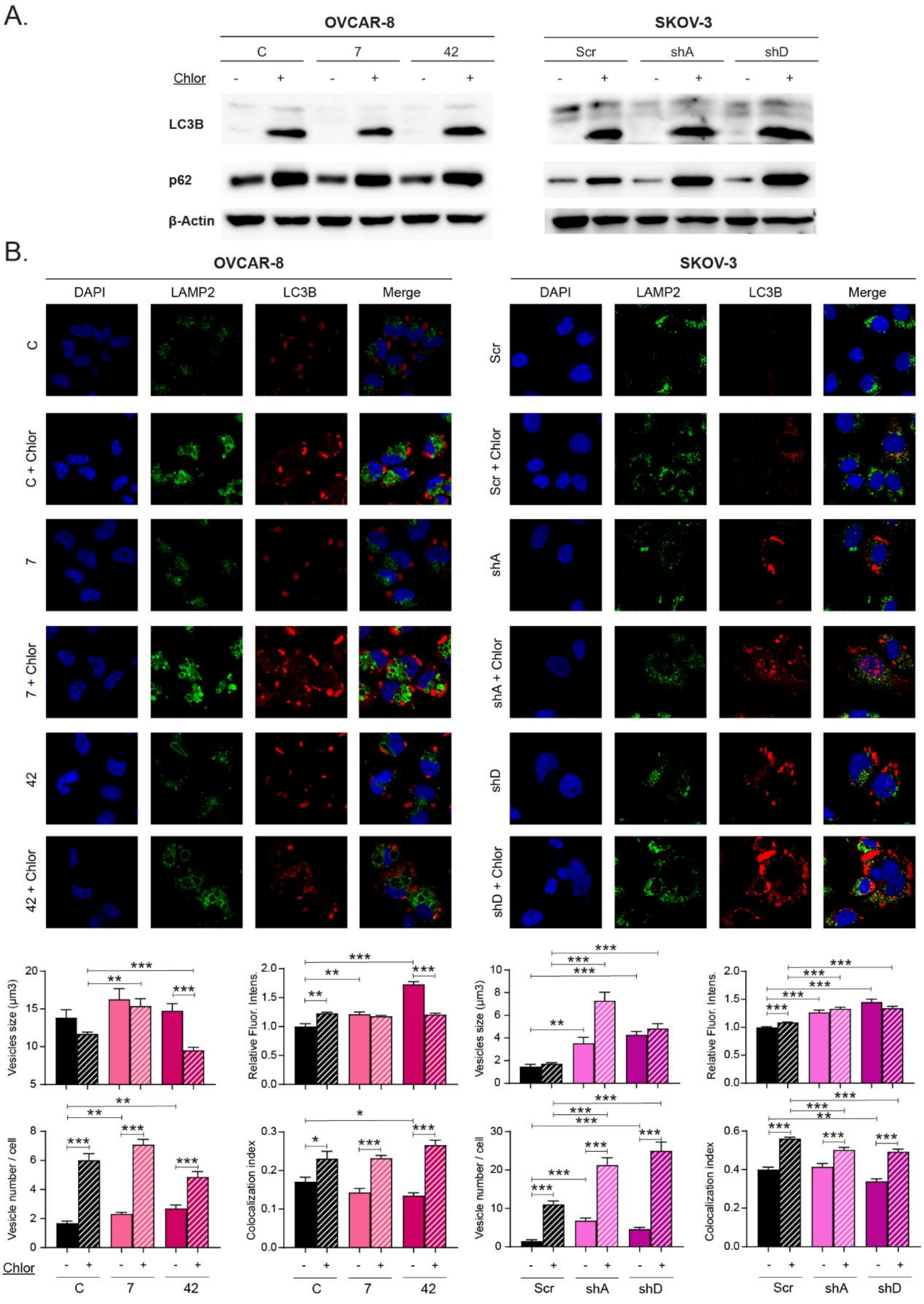

**SUPPLEMENTARY FIGURE 4**

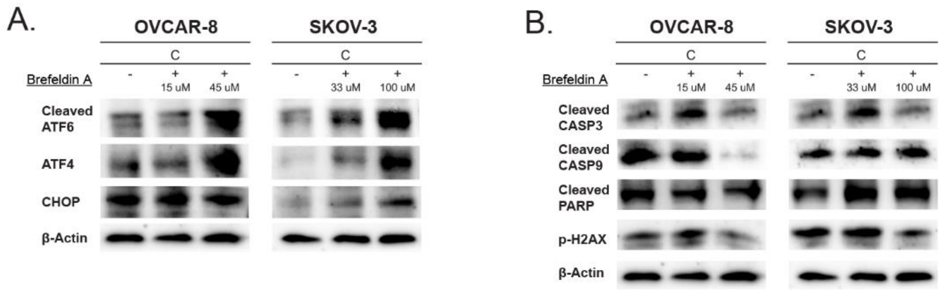

### SUPPLEMENTARY FIGURES LEGENDS

**Supplementary Figure 1. Validation of the HOOK2 reduction models generated in ovarian cancer cell lines OVCAR-8 and SKOV-3. (A)** WB analysis of HOOK2 expression in HOOK2 CRISPR clones within OVCAR-8 and SKOV-3 cell lines. Clones selected for use in the OVCAR-8 line are highlighted in red. **(B)** DNA sequencing of the sgRNA region in the selected CRISPR clones in the OVCAR-8 line, compared to the original sequence. **(C)** q-RT-PCR of HOOK2 expression in selected CRISPR clones in the OVCAR-8 line and selected shRNAs in the SKOV-3 line. The mean of 3 independent experiments ± SEM is presented. Statistical analysis was performed with Student's t test (*$p < 0.05$; **$p < 0.01$; ***$p < 0.001$). The lack of an asterisk indicates that the data do not reach statistical significance.

**Supplementary Figure 2. Ontology term analysis of proteins altered by the downregulation of HOOK2 levels.** Ontology term analysis of **(A)** upregulated and **(B)** downregulated proteins upon HOOK2 reduction performed in the ShinyGO 0.80 web resource. Breakdown of proteins **(C)** upregulated and **(D)** downregulated upon HOOK2 reduction implicated in the processes depicted in Figure 2B and C. **(E)** Heatmap of relative normalized spectral counts of proteins involved in protein folding and response to ER stress in cells with downregulated HOOK2.

**Supplementary Figure 3. Validation of chloroquine treatment. (A)** Protein levels of autophagy-associated proteins in HOOK2-downregulated cells treated with chloroquine. **(B)** Immunofluorescence staining of LAMP2 (in green) and LC3B (in red) proteins in HOOK2-downregulated cells treated with chloroquine. DAPI (in blue) was utilized for nuclear staining, and the merge of the three markers is presented. Autophagosome size, number, relative fluorescence intensity, and the colocalization index of autophagosomes-lysosomes were quantified using ImageJ software. Analysis involved a minimum of 100 cells for each condition. Statistical analysis was performed with Student's t test (*$p < 0.05$; **$p < 0.01$; ***$p < 0.001$). The lack of an asterisk indicates that the data do not reach statistical significance.

**Supplementary Figure 4**. **Validation of brefeldin A treatment.** Protein levels of **(A)** UPR-associated proteins and **(B)** apoptosis-associated proteins in ovarian cancer cells treated with different doses of the ER stress inducer brefeldin A.
